# Azacytidine restores T cell function in AML by modulating DNA methylation

**DOI:** 10.64898/2026.06.14.732148

**Authors:** Ravina Pandita, Yoko Kosaka, Jessica S. Mulkey, Cora E. Layman, Brett A. Davis, Lucia Carbone, Evan F. Lind

**Affiliations:** Graduate Program in Biomedical Sciences, Oregon Health & Science University, Portland Oregon, USA; Department of Molecular Microbiology and Immunology, Oregon Health & Science University, Portland Oregon, USA; Knight Cancer Institute, Oregon Health & Science University, Portland Oregon, USA; Knight Cardiovascular Institute, Oregon Health & Science University, Portland, OR, USA; Department of Molecular and Medical Genetics, Oregon Health & Science University, Portland, OR, USA; Division of Genetics, Oregon National Primate Research Center, Beaverton, OR, USA; Department of Cell, Development and Cancer Biology. Oregon Health & Science University, Portland, OR, USA; Department of Pediatrics, Oregon Health & Science University, Portland, OR, USA

**Author notes:** Corresponding author: Evan F Lind.

## Abstract

AML is an aggressive blood cancer associated with poor clinical outcomes. Chemotherapy remains the standard of treatment, but unfortunately relapse is very common, highlighting the need for alternative therapies. T cell dysfunction and exhaustion are prominent in AML and may represent a barrier to effective immunotherapy yet remains poorly studied in AML. DNA methylation is a major driver of T cell exhaustion and inhibition of *de novo* methylation can block exhaustion and restore T cell function in chronic viral infections and other cancers but is understudied in AML. Here, we investigated the impact of azacytidine (Aza), an FDA-approved hypomethylating agent, on T cell exhaustion in AML. Using a spontaneous AML mouse model and samples from patients with AML, we found that Aza treatment modulates T cell function. *In vivo* Aza-treatment of AML-bearing mice decreased tumor burden and reshaped CD8+ T cell states, with increases in frequencies of memory subsets and decreases in regulatory T cells (Tregs). Functionally, Aza treatment overcame the impaired proliferation displayed by both CD4 and CD8+ T cells in our model. DNA methylation sequencing of T cells after Aza treatment revealed hypomethylation and increased expression of stem-like precursor gene *TCF7* and *E2F2*, a regulator of cell cycle progression and proliferation. Similar changes in phenotypes were observed in cultures of AML patient samples treated with Aza. Collectively, we show that Aza remodels epigenetic and functional states in AML and has the potential to reverse T cell exhaustion, with enhanced memory and proliferation capacity. Our work generates a mechanistic framework that provides rationale of combining hypomethylating agents with T cell-based immunotherapies in this lethal disease.

**Data Sharing Statement:** RRBS data is available in GEO under the accession number GSE328721. For original data please contact Dr. Evan F. Lind.

**Key Points:** Azacytidine mediated epigenetic modulation can alleviate T cell exhaustion in AML

**Translational Relevance:** Immune therapy has shown limited efficacy in AML, despite increasing evidence of T cell dysfunction in this malignancy. Azacytidine (Aza) is an FDA approved drug for AML, but patients develop therapy resistance and relapse. Studies have mainly focused on Aza’s tumor intrinsic effects. In this study, we investigated the impact of Aza on immune function, especially T cell exhaustion in AML, since exhaustion is a major mechanism of disease resistance. We demonstrated that Aza can modulate T cell phenotype and restore T cell proliferation. Mechanistically, Aza induces epigenetic reprogramming in T cells and increases the expression of a stem-like precursor marker, TCF7. By shifting the focus on T cell biology, our study provides a rationale for combining Aza with other immunotherapies that can enhance durable immune responses in this malignancy.

## Introduction

Acute myeloid leukemia (AML) is a highly aggressive disease with a 5-year survival rate of approximately 33%^1^. While intensive chemotherapy has been the standard of care over the last five decades, many patients develop therapy resistance and relapse. The only curative therapy remains allogeneic stem cell transplant which is effective in great part due to graft vs leukemia immunity^2–4^. The success of stem cell transplantation highlights the potential efficacy of immune-based therapies in AML^5^. Consistent with this, we and others have shown that immune suppression, especially T cell dysfunction is a prominent feature in patients with AML at the time of diagnosis^6–9^.

A key mechanism driving T cell dysfunction in chronic viral infections and cancer is T cell exhaustion, which is also observed in AML^8,10–13^. Our prior work demonstrated that T cells in the bone marrow of a subset of newly diagnosed patients with AML were dysfunctional^6^. However, checkpoint blockade was able to restore T cell function *in vitro* in a majority of these samples. In the exhaustion hierarchy, the ability of T cells to respond to checkpoint blockade is largely retained within a subset of precursor exhausted T cells (TPex), which maintain a stem-like progenitor state^14^. A defining feature of these TPex cells is the expression of the transcription factor TCF1, which is essential to retain effector function and responsiveness to checkpoint blockade^15–17^.

Epigenetic remodeling is a fundamental driver of T cell exhaustion^18,19^. Exhausted T cells adopt a distinct epigenetic state compared to effector or memory subsets^20^. *De novo* DNA methylation by DNMT3a can cause T cell exhaustion in chronic viral infection settings^21^. Notably, genetic or pharmacologic disruption of DNA methylation can restore T cell function, and enhance immunotherapy responses in several cancers^22,23^.

Aberrant DNA methylation is observed in a subset of patients with AML, especially those harboring mutations in epigenetic regulators, such as TET2, IDH1/2 and DNMT3A^24–27^. These epigenetic alterations provide a rationale for the use of hypomethylating agents such as azacytidine (Aza), which is an FDA-approved therapy for AML and myelodysplastic syndromes, commonly used for patients that are ineligible for intensive chemotherapy or hematopoietic stem cell transplants^28^.

Although 30%-40% of patients initially respond to therapy with Aza, patients eventually relapse, suggesting resistance mechanisms. While Aza has historically been studied for its direct effect on the tumor, emerging evidence suggests that Aza can also modify immune function^29–31^. However, how Aza impacts T cell exhaustion and epigenetic reprogramming in the setting of AML remains poorly described.

Here, we use a spontaneous, genetically engineered mouse model of AML developed in our lab that harbors FLT3-ITD and loss of TET2, limited to the myeloid compartment. This AML mouse model recapitulates leukemia burden and features of T cell exhaustion, including the loss of T cell proliferative capacity^7,32^. We show that Aza treatment reduces tumor burden, reshapes CD8+ T cell memory subsets, and restores T cell proliferation. Reduced Representation Bisulfite DNA Sequencing of T cells after *in vivo* Aza treatment shows epigenetic remodeling of transcription programs associated with stem-like exhausted state. These findings are also observed in samples from patients with AML, supporting the role of Aza as an immune modulating agent that can also impact T cell exhaustion programs in AML.

## Methods

### AML model

All mice in this experiment were handled according to the guidelines provided by an approved Institutional Animal Care and Use Committee (IACUC) protocol IP00000907. Mice were purchased from Jackson labs and were housed in pathogen free conditions.

Our mice with AML harbor an internal tandem duplication of FLT-3 (FLT3-ITD)^33^ and a TET2^34^ deletion which is restricted to the myeloid compartment, leaving the lymphoid compartment fully intact. Mice with FLT3-ITD have an endogenous FLT3 promoter (B6.129-*Flt3tm1Dgg*/J, stock 011112), which is then crossed with mice having floxed sites for TET2, TET2f (strain B6;129S-*Tet2tm1.1Iaai*/J, stock 017573). The mice generated from this cross are then crossed with LysM-Cre mice (B6.129P2- *Lyz2tm1(cre)Ifo*/J, stock 004781). The mice used in this experiment were heterozygous for FLT-3-ITD and LysM-Cre, and homozygous for TET2f. Mice were aged between 15-55 weeks of age and were randomly assigned into treatment groups after leukemia burden was confirmed. Sample size was determined based on preliminary experiments, which informed expected effect size and was selected to ensure appropriate statistical power. Both male and female mice were used in the study and no blinding was performed.

### Aza treatment and *in vivo* experimental design

Azacytidine was purchased from Millipore Sigma (PHR1911). Fresh drug was prepared daily in PBS immediately prior to administration. Dosing regimen is shown in **Figure 1A**. This dosing schedule was determined based on prior optimization experiments performed in our lab^35^.

**Figure 1:**
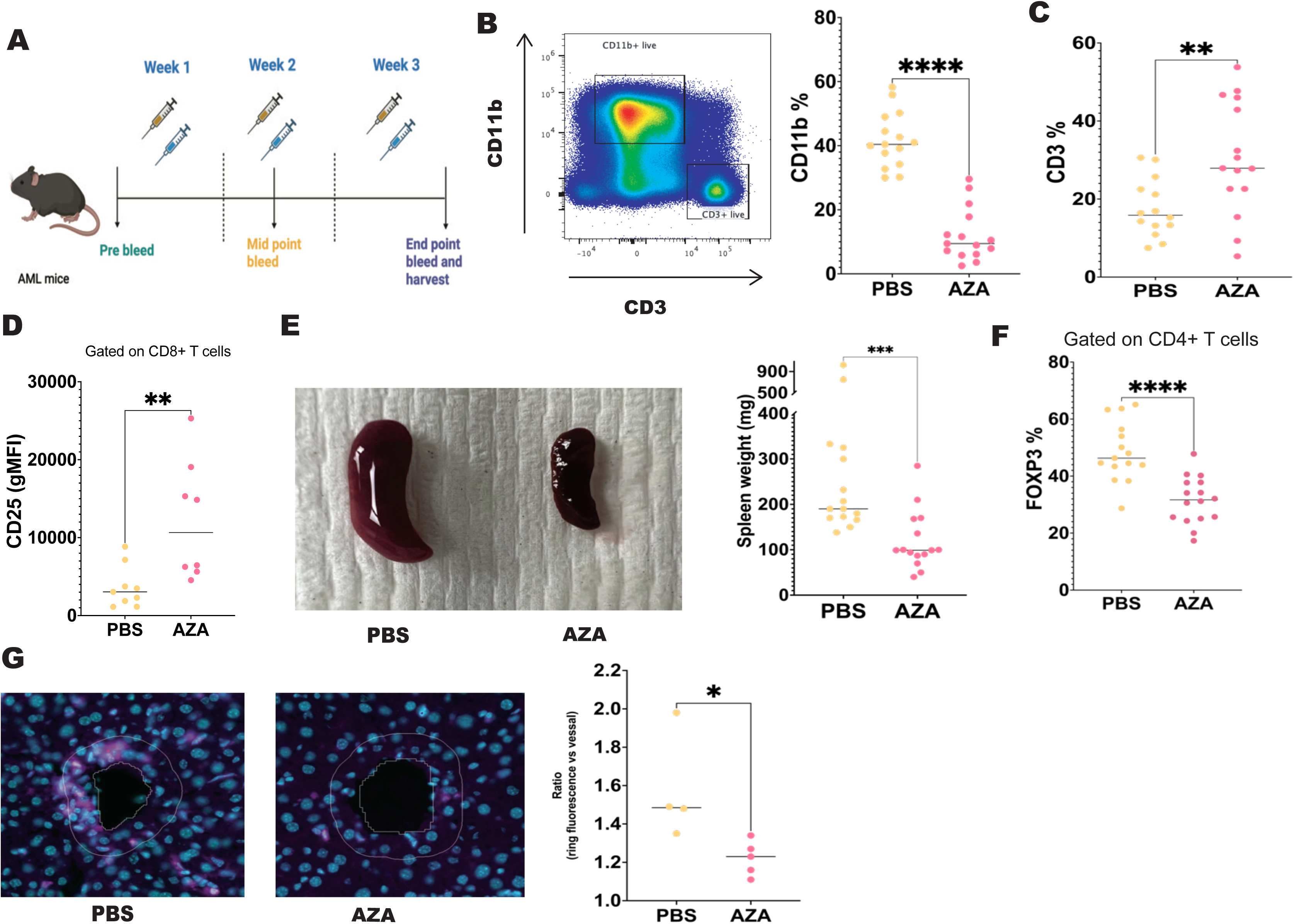
Impact of Aza on tumor burden in AML mouse model. (A) Experimental schematics of *in vivo* azacytidine (Aza) treatment schedule. FLT3-ITD TET2-/-leukemic mice received Aza (1mg/kg, intraperitoneal) or vehicle control (PBS) for 5 consecutive days followed by a 2-day break, repeated over three weeks. Peripheral blood was collected to assess tumor burden and T cell phenotype. Mice were randomized into treatment groups by measuring leukemia burden (% CD11b+ cells in the blood) prior to treatment. Mice were euthanized at the study endpoint (3 weeks) for a comprehensive analysis of blood, spleen and liver. (B) Representative flow cytometry dot plot and quantification of CD11b+ cells among live splenocytes from PBS and Aza-treated mice after three weeks of treatment (p-value= <0.0001). Cells were gated on live singlets. (C) Frequency of CD3+ T cells among live splenocytes from PBS and Aza-treated mice assessed by flow cytometry (p-value= 0.0082). (D) T cell activation following three weeks of *in vivo* treatment, assessed by flow cytometry. Geometric mean fluorescence intensity (gMFI) of CD25 expression was measured on CD8+ T cells isolated from blood of PBS and Aza-treated mice (p-value= 0.0055). (E) Representative photographic images of spleens (left) and spleen weights (right) harvested from PBS and Aza-treated mice. (F) Frequency of regulatory T cells (CD4+ FOXP3+) among CD4+ T cells from spleens of PBS and Aza-treated mice, as assessed by flow cytometry (p-value= <0.0001). (G) Immunofluorescence imaging analysis of liver sections from PBS and Aza treated mice. Based on DAPI nuclear staining signal, 15-25 blood vessels were randomly selected as objects per tissue section within the cross-section of the vessel by magic wand tool in ZEISS Arivis Pro 4.4.0. The proximity of myeloid cells to vessels was quantified as a ratio of mean signal intensity within a 25uM outward expanded ring to the mean intensity in the vessel volume. Representative images show CD11b+ myeloid cells in and around blood vessels, where pink is CD11b+ cells and light blue is DAPI. Quantification shows the ratio of CD11b fluorescence intensity in the ring compared to vessels, measuring average intensity from positive cells (p-value= 0.0159). Data are pooled from 1-4 independent experiments, with n= 4-13 mice per group, as shown. Statistical analyses were performed using Mann-Whitney U test. p-values as indicated.

### Peripheral blood analysis

For automated blood counts, 20mL of peripheral blood was collected into EDTA-coated tubes and analyzed using an automated hematology analyzer (Element HT5). Data was collected on HenskaView software.

### Flow cytometry

Spleens were mechanically disassociated, and RBCs were lysed using ACK buffer. Single cell suspensions at 1-5 million cells per sample were used for flow cytometry staining. Cells were first incubated in Zombie Aqua viability dye (BioLegend). Cells were then washed in flow buffer (2% calf serum, 0.5% sodium azide, PBS) twice before incubating with Fc receptor blocking reagent (Human or Mouse TruStain FcX, Biolegend). Surface antibodies **(Table 1/2)** were then added, and samples were incubated for 30 minutes.

**Table 1:**
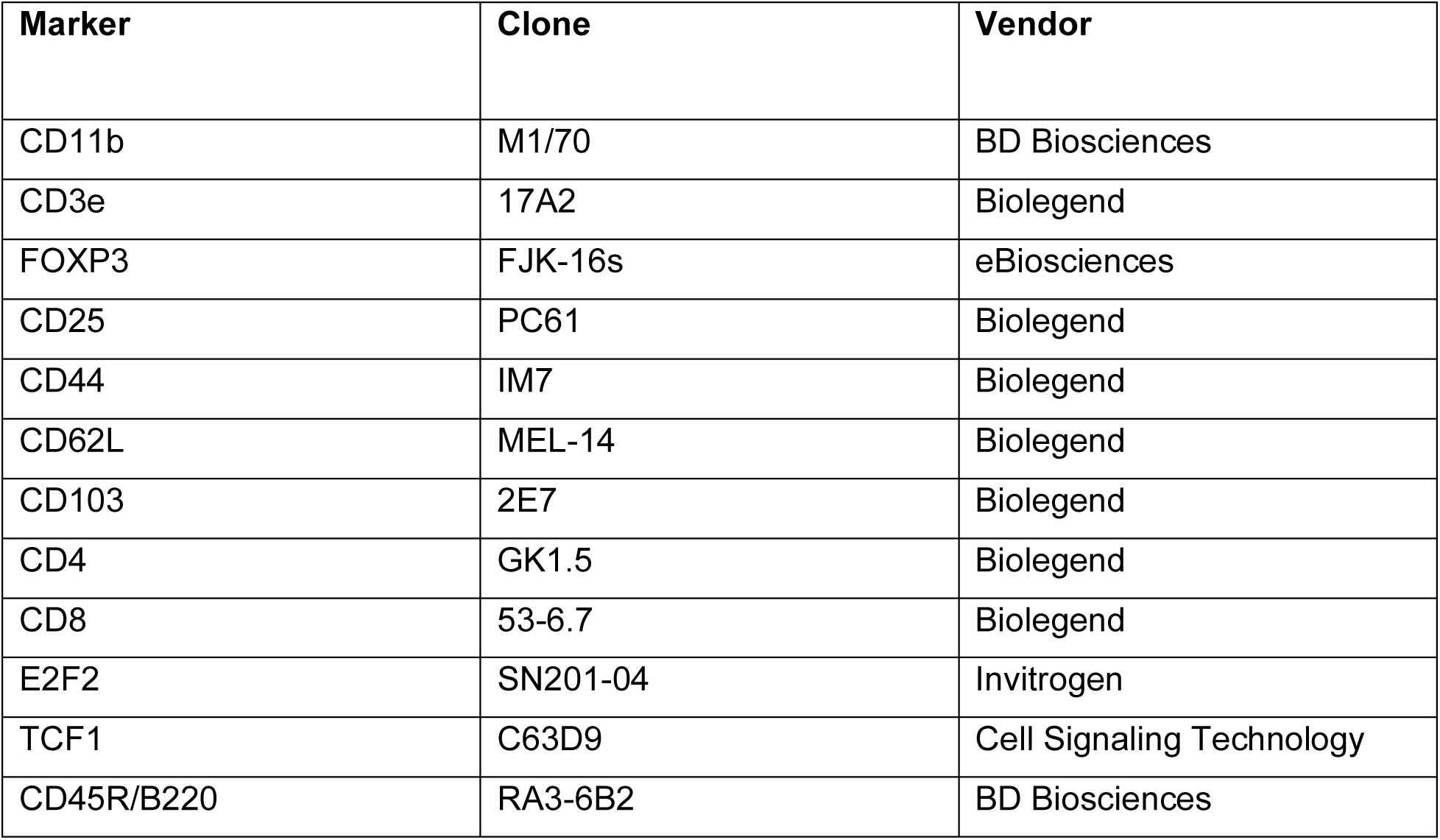
Mouse flow cytometry antibodies-.

**Table 2:** Human flow cytometry antibodies-.

| Marker | Clone | Vendor |
| --- | --- | --- |
| CD33 | WM53 | Biolegend |
| CD3 | HIT3a | Biolegend |
| FOXP3 | 206D | Biolegend |
| CD25 | BC96 | Biolegend |
| TCF1 | C63D9 | Cell Signaling Technology |
| E2F2 | SN201-04 | Invitrogen |
| CD103 | B-Ly7 | eBiosciences |
| CD4 | RPA-T4 | Biolegend |
| CD8 | RPA-T8 | Biolegend |

For intracellular staining, cells were fixed and permeabilized using Foxp3/Transcription Factor Staining Buffer Set (eBiosciences). Fc receptor blocking was performed again. Cells were subsequently incubated with intracellular antibody cocktail **(Table 1/2)** overnight at 4°C. Samples were then washed with 1X permeabilization buffer, resuspended in flow buffer and were acquired on Cytek Aurora. Data analysis was performed using FlowJo software (version 10.10.0). Gating strategies are shown in **Supplemental figure 3D** and relevant panels.

### Mouse *ex vivo* T cell proliferation assay

Splenocytes were harvested and cells were labelled with 2mM Carboxyfluorescein Succinimidyl Ester (CFSE) (Life Technologies, Thermo Fisher). 96 well round bottom plates were pre-coated with 1-2.5 mg/ml hamster IgG (clone: HTK888, Biolegend) or anti-CD3 antibody (clone: 145-2C11, Biolegend). CFSE-labeled cells were then plated and cultured for 72 hours at 37°C. Following incubation, cells were harvested, stained for surface markers and analyzed using flow cytometry to assess proliferation based on CFSE dilution.

### *Ex vivo* experiments using patient samples with AML

All patients provided informed consent and were approved under IRB protocol #00004422, “Pathogenesis of Acute Leukemia, Lymphoproliferative Disorder and Myeloproliferative Disorders” (PI: Marc Loriaux, MD, PhD). Frozen peripheral blood mononuclear cells (PBMCs) from patients with AML were thawed, washed and resuspended in IMDM with 20% FBS supplemented with IL-3 (10ng/ml), IL-6 (100ng/mL), SCF (100ng/mL), IL-2 (200U/mL), IL-7 (5ng/mL) and IL-15 (5ng/mL). Cells were plated in 96-well round bottom plates, coated with 1mg/mL mIgG (clone: MG2b-57, BioLegend) or anti-CD3 (clone: UCHT1, BioLegend) and treated daily with Aza (500nM or 1mM) or vehicle (PBS, in mL accordingly) for 5 days. Cells were then harvested and processed for flow cytometry as described above.

### T cell sorting

Splenocytes were stained with antibodies against CD4 and CD8 and sorted using BD Symphony S6. Sorted CD4+ and CD8+ T cells were collected into flow buffer (without sodium azide) for further downstream epigenetic analysis.

### DNA isolation and Reduced Representation Bisulfite Sequencing (RRBS)

DNA isolation was performed on sorted T cells using Puregene Cell kit (Qiagen) according to manufacturer’s guidelines. Zymo Clean and Concentrator Kit was used to clean up the collected DNA. After enriching for CpG dense regions, library preparation was performed using NEBNext Ultra II Modules (New England Biolabs) using methylated adaptors. Next, the EZ DNA Methylation-Gold Kit (Zymo Research) was used to perform bisulfite conversion. PCR amplification of DNA was performed using Q5U polymerase and NEBNext Multiplex Oligos for Illumina (New England Biolabs). After quantifying the libraries, they were normalized and sequenced at the OHSU Massively Parallel Sequencing Shared Resource (MPSSR) sequencing core.

### RRBS data analysis

Raw sequences were assessed for quality with FastQC (v0.11.9)^36^ and trimmed for quality and adapters with TrimGalore (v0.6.7)^37^ using the –rrbs parameter. Trimmed reads were aligned to the mouse reference genome from Ensembl, GRCm38, with Bismark (v0.24.1)^38^ using default parameters. The bismark_methylation_extractor script was used to obtain coverage files for downstream analysis. The R library Methylkit^39^ was used to perform differential methylation analysis by tiling the genome into 1kb non-overlapping regions and averaging the CpG methylation rates within each region. Logistic regression was applied, and a significance cut-off of q-value < 0.1 and absolute value of the minimum methylation percent difference >10% was applied to identify significant differentially methylated regions (DMRs). The ChIPseeker^40^ library were used with Ensembl annotation (GRCm38.99) to annotate DMRs to overlapping and nearby genes. Pathway enrichment analysis was performed on the DMRs using Genomic Regions Enrichment of Annotations Tool (GREAT)^41,42^. For the CpG plots, CpG methylation was obtained over regions of interest from the bedgraph files produced by Bismark methylation extractor. Percent methylation values were averaged over group replicates. Gene models were obtained from Ensembl annotation for the mouse reference genome GRCm38 (GRCm38.99 annotation). This data is available in GEO, and the accession number is **GSE328721**.

### Immunofluorescence sample preparation and imaging

Livers were harvested and fixed in 4% PFA overnight at 4°C. Following day, tissues were washed in PBS and cryoprotected using 30% sucrose (99%, Thermo scientific) overnight. Tissues were embedded in OCT compound (Fisher Scientific) and were frozen rapidly using isopentane cooled with dry ice. Tissues were sectioned using a cryostat (Leica CM 3050s) and mounted on glass slides. Samples were then blocked using 2% rat serum (Invitrogen) in a humidified chamber for 1 hour. Sections were incubated with CD11b antibody for 1 hour, followed by Vector TrueView autofluorescence quencher kit (Vector Laboratories). DAPI was used for nuclear staining. Samples were mounted using VECTASHIELD Vibrance Antifade Mounting Medium (Vector Laboratories).

For image acquisition, Hamamatsu Orca Flash monochrome camera was used along with ZEISS Axioscan 7 slide scanner with a Plan-Apochromat 20x/0.8 objective (0.324 µm/pixel resolution).

### Statistical analyses and figures

Statistical analysis was performed using GraphPad PRISM 10. Mann-Whitney U test, paired-t test or repeated measures ANOVA was performed to compare the two treatment groups, as appropriate. A p-value of <0.05 were considered statistically significant. All data is presented as Mean ± SD for bar graphs. For dot plots, each dot represents an individual mouse, and the horizontal line indicates the median. Schematics in the figures were created using Biorender.com. Adobe illustrator was used to create figures.

## Results

### Azacytidine treatment reduces leukemia burden and alters lymphocyte populations in mice with AML

Mice were treated with Aza or vehicle (PBS) for 3 weeks with treatment regimen as outlined **(Figure 1A)**. After three weeks, Aza treatment led to a significant reduction in leukemia burden, as evidenced by a decreasing frequency of CD11b+ myeloid cells in both spleen and blood **(Figure 1B & Supplemental figure 1A)**. Aza-treated mice displayed an increased frequency of CD3+ T cells in the spleen compared to vehicle controls **(Figure 1C)**. The changes in T cell frequency were also tissue-dependent and limited to the spleen **(Supplemental figure 1B).** Automated blood counts revealed a shift in circulating immune populations following Aza treatment, including a decreased monocyte frequency and an increase in lymphocytes **(Supplemental figure 1C, D)**. Consistent with this, Aza-treated mice showed increased expression of the activation marker CD25 on CD8+ T cells **(Figure 1D)** and on CD4+ T cells **(Supplemental Figure 1E)** in blood, further supporting enhanced T cell responsiveness following Aza treatment. This change was not significant in T cells from spleen **(Supplemental Figure 2F-G)**, again suggesting that these effects are tissue-specific in AML. Additionally, Aza treatment led to a marked reduction in splenomegaly as shown by representative images and quantification of spleen weight **(Figure 1E)**. Furthermore, there was a decrease in frequency of regulatory T cells (Tregs) after Aza treatment in both spleens and blood **(Figure 1F & Supplemental figure 1H)**. We also identified B cells (by B220), which was significantly higher in frequency in the spleens of Aza-treated mice **(Supplemental figure 1 I**). Similar to T cell changes, increased B cell populations were localized to the spleen **(Supplemental figure 1 J)**. We have previously observed tumor infiltration in the livers of animals with AML^32^. We performed immunofluorescence imaging of CD11b on liver sections from these mice and found reduced myeloid infiltration around blood vessels in the Aza treatment group compared to vehicle mice **(Figure 1G)**. Overall, Aza treatment reduced tumor burden in our AML mouse model.

### Aza treatment *in vivo* restores the proliferation capacity of T cells in animals with AML

T cells in our AML mouse model have a defect in the ability to proliferate in response to stimulation with anti-CD3 *ex vivo*^7,32^. To determine whether Aza has an impact on T cell function, we conducted *ex vivo* proliferation assays using splenocytes harvested after three weeks of Aza treatment. Cells were labeled with CFSE and cultured on irrelevant IgG or anti-CD3 coated plates for three days **(Figure 2A)**.

**Figure 2:**
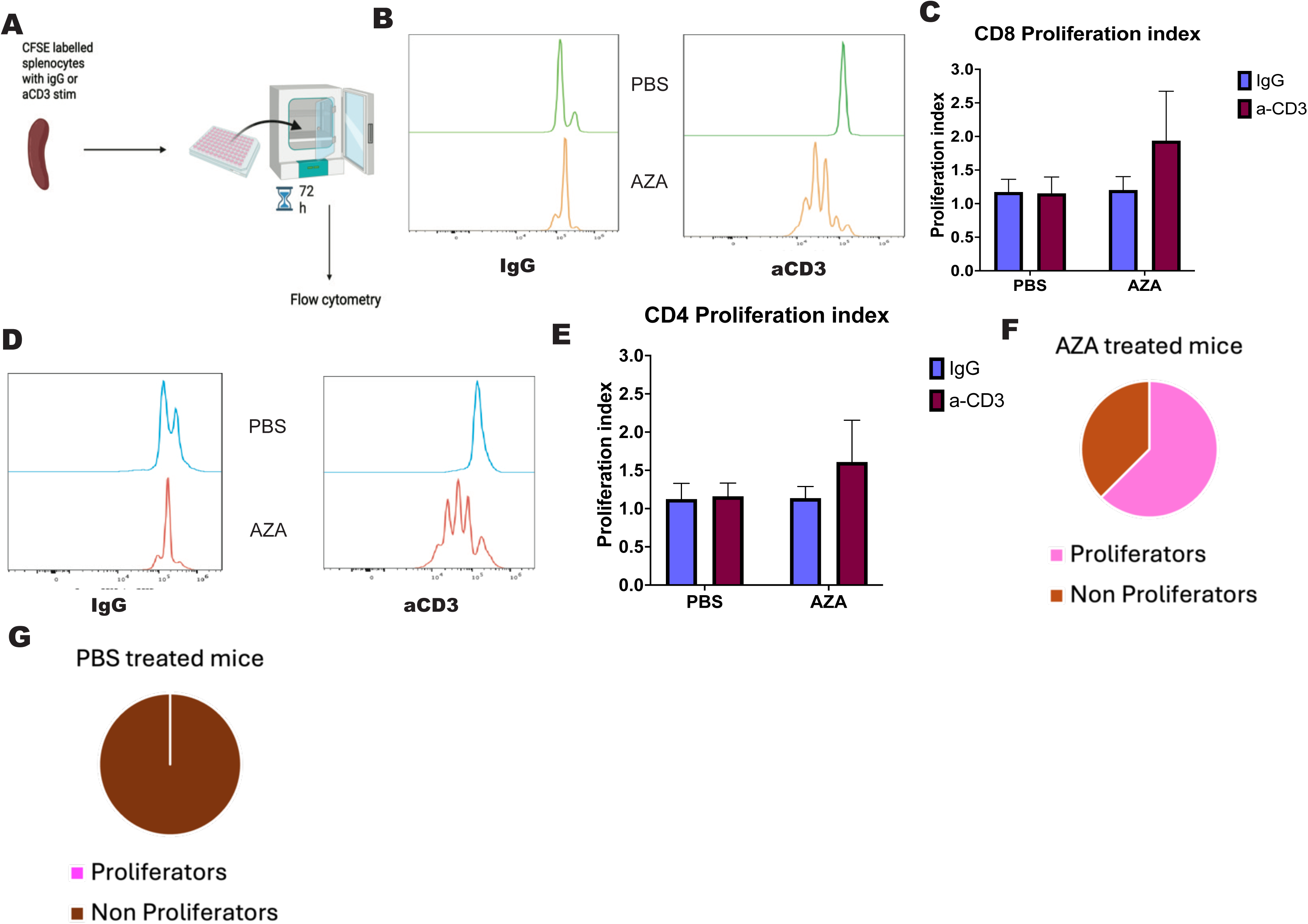
T cell proliferation following Aza treatment. (A) Schematic of the *ex vivo* T cell proliferation assay. Splenocytes were harvested from leukemic mice following three weeks of *in vivo* treatment with Aza or PBS. Cells were labelled with CFSE and cultured on IgG or anti-CD3-coated plates for 72 hours prior to flow cytometric analysis. (B) Representative CFSE histograms showing proliferation profiles of CD8+ T cells under IgG or anti-CD3 stimulation conditions from PBS or Aza treated mice. Cells were gated on CD3+ T cells, and subsequently on CD8+ T cells prior to CFSE analysis. (C) Proliferation index quantification by the FlowJo proliferation tool for CD8+ T cells, with IgG and anti-CD3 stimulation conditions from PBS or Aza-treated groups. (D) Representative CFSE histograms from CD4+ T cells in both treatment groups, on IgG or anti-CD3 stim. (E) Proliferation index quantification by the FlowJo proliferation tool for CD4+ T cells, with IgG and anti-CD3 stimulation conditions from PBS or Aza-treated groups. (F) Pie chart summarizing the proportion of Aza-treated mice exhibiting measurable proliferation/CFSE dilution in response to anti-CD3 stimulation across experiments. (G) Pie chart summarizing proliferative responses of PBS treated mice under anti-CD3 stimulation conditions across experiments. Data is pooled from two independent experiments with n=8-9 mice per treatment group, as shown. Statistical analyses were performed using Mann-Whitney U test. p-values as shown.

As previously observed, our negative control IgG did not induce proliferation in either group. Upon anti-CD3 stimulation, T cells from PBS-treated mice failed to proliferate, whereas both CD4+ and CD8+ T cells from Aza treated mice exhibited robust CFSE dilution, indicating increased proliferative capacity compared to T cells in the PBS-treated animals. This can be visualized by representative CFSE dilution histograms, proliferation index measurements in CD8+ **(Figure 2B-C)** and CD4+ gated T cells **(Figure 2D-E)**. This can also be visualized in the flow cytometry plots **(Supplemental Figure 2A)**. Approximately two-thirds of Aza-treated mice demonstrated measurable T cell proliferation, while we observed no proliferation of T cells from the PBS-treated mice **(Figure 2F-G)**, suggesting Aza can restore T cell function.

### Aza alters T cell memory distribution

Given prior studies linking DNA methylation to T cell differentiation and memory fate, we assessed the impact of Aza on CD8+ T cell memory subsets. Using CD44 and CD62L expression to define memory populations **(Figure 3A)**, we observed a significant increase in the proportion of central memory CD8+ T cells in the peripheral blood and spleens of Aza-treated mice **(Figure 3B & Supplemental figure 3A)**.

**Figure 3:**
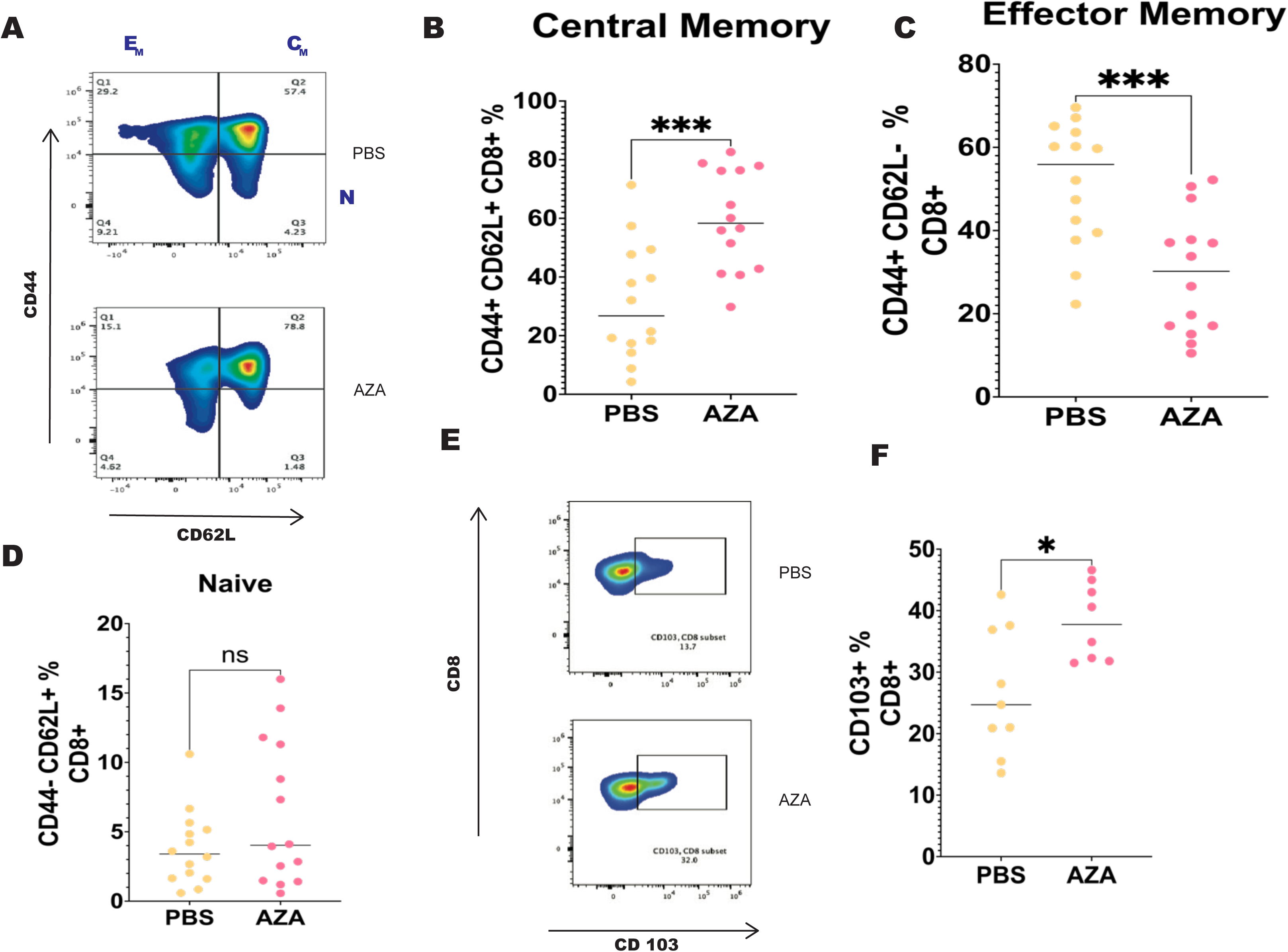
Memory phenotype distribution of CD8+ T cells following Aza treatment. (A) Representative flow cytometry dot plots showing the gating strategy used to define CD8+ T cell memory subsets based on CD44 and CD62L expression in blood of CD8+ T cells from PBS and Aza-treated mice. It also shows a representative comparison between the two groups. E_M_ is effector memory, C_M_ is central memory and N is naïve. (B-D) Frequencies of (B) central memory (CD44+ CD62L+) (p-value= 0.0006), (C) effector memory (CD44-CD62L+) (p-value= 0.0009), and (D) naïve (CD62L-CD44-) (p-value= 0.4544), CD8+ T cell subsets in the peripheral blood following three weeks of treatment. (E, F) Representative flow cytometry plots showing CD103 expression on CD8+ T cells (E) and quantification of CD103+ CD8+ T cell frequency among total CD8+ T cells (F) (p-value= 0.0274) in spleens following three weeks of PBS or Aza treatment. Data is pooled from 2-4 independent experiments with n=8-14 mice per group, as indicated. Statistical analyses were performed using Mann-Whitney U test, and p-values are mentioned.

Conversely, Aza treatment resulted in a significant reduction in effector memory CD8+ T cells **(Figure 3C & Supplemental figure 3B)**, consistent with findings from viral infection models and CAR-T cells, where inhibition of DNA methylation programs promote memory over terminal effector differentiation^23,43^. No significant changes were observed in naïve populations **(Figure 3D & Supplemental figure 3C)**. These data suggest that Aza shifts CD8+ T cells towards a less differentiated, more stem-like memory biased state.

Tissue-resident memory cells have been associated with positive outcomes in solid tumors and as a marker of response to immune checkpoint therapy. A recent study demonstrated that patients with leukemia that attained complete remission after chemotherapy had higher frequencies of CD103+ T cells as compared to *de novo* or relapsed/refractory patients implying that this subset was representative of possible anti-tumor immunity^44^. For this reason, we examined tissue-resident memory populations based on the expression of CD103. We found that Aza-treated mice have a significant increase in the frequency of CD103+ CD8+ T cells in the spleen **(Figure 3E-F)**.

### Aza induces epigenetic remodeling of T cells

Next, we wanted to evaluate changes that Aza imparts on the DNA methylation landscape of T cells in AML. We performed reduced representation bisulfite sequencing (RRBS) on sorted CD4+ and CD8+ T cells from spleens of these mice after treatment **(Figure 4A)**.

**Figure 4:**
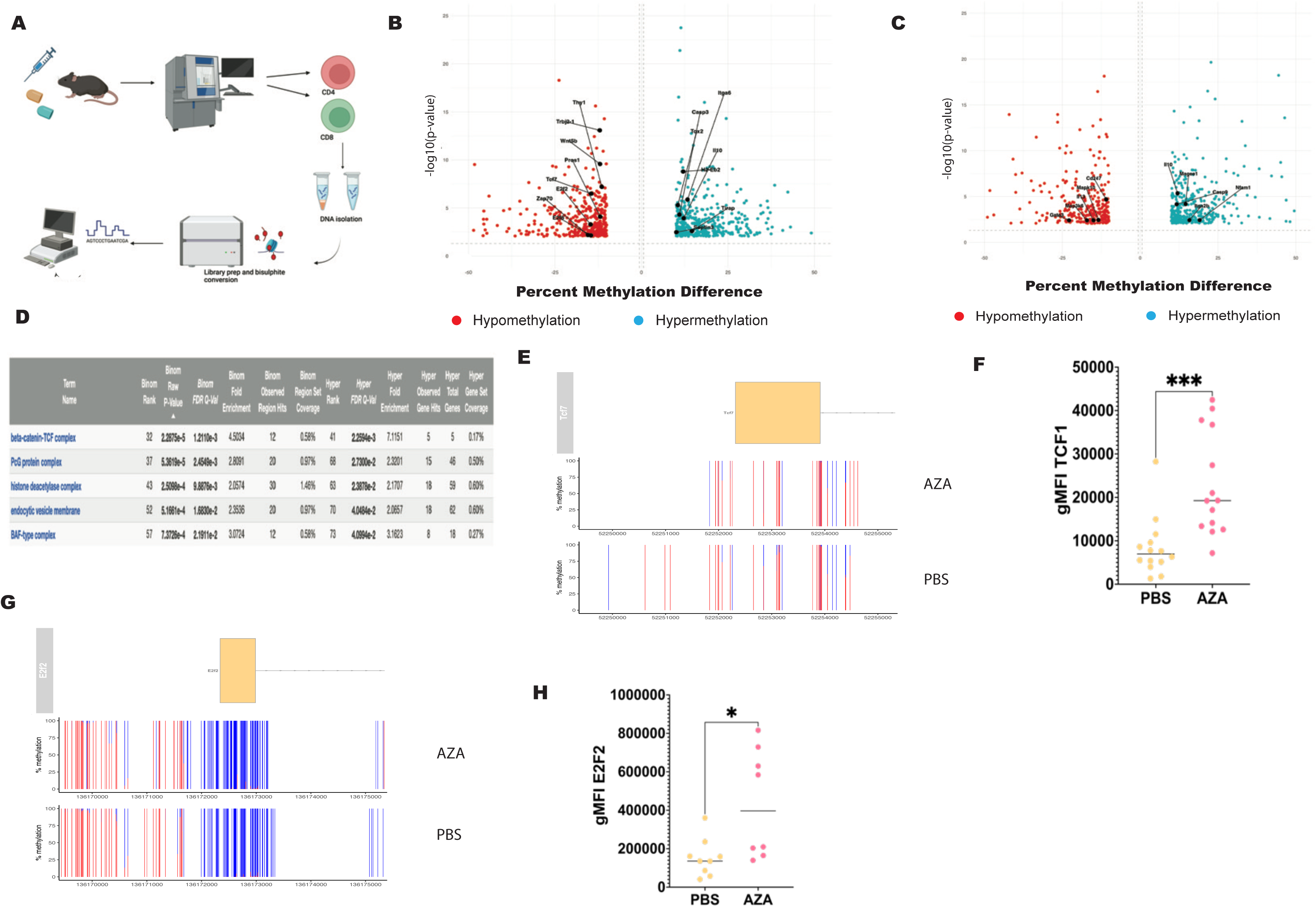
Reduced representation bisulfite sequencing (RRBS) of CD4+ and CD8+ T cells following Aza treatment. (A) Schematic of the experimental workflow for RRBS. Following three weeks of *in vivo* treatment of leukemic mice with Aza or PBS, CD4+ and CD8+ T cells were isolated from spleens by flow sorting. Genomic DNA was extracted from sorted populations and processed for RRBS library preparation and bisulfite sequencing, followed by differential methylation analysis. (B, C) Volcano plots displaying differentially methylation analysis of manually highlighted immune-related genes in CD8+ (B) and CD4+ (C) T cells, showing percent difference methylation on a log (10) scale. A methyl percent difference of > 10% was used as a cut-off and the plot was generated in R-studio. All DMRs plotted were located in the promoter regions. (D) Table indicating the top 5 enriched pathways identified by GREAT analysis using DMRs that were hypo- or hyper-methylated in CD8+ T cells following Aza treatment. Pathway statistics were generated using BED files and the GREAT tool. (E) Group averages of CpG percent methylation levels around the transcription start site (TSS) of the *TCF7* locus in CD8+ T cells from PBS and Aza-treated mice. Each bar represents a covered CpG. The ratio of red to blue represents the ratio of methylated to unmethylated reads, respectively. RRBS analysis was performed using methylkit with defined cutoffs for differential methylation as described in methods. The R libraries Gviz and ggplot2 were used to plot the gene model and methylation data. (F) Flow cytometric analysis of TCF1 (p-value = 0.0001) expression on CD8+ T cells from the blood of PBS or Aza treated mice. Graph shows quantification of gMFI after gating on live CD3+ CD8+ T cells. (G) Group averages of percent CpG methylation around the TSS of the *E2f2* locus in CD8+ T cells from PBS or Aza-treated mice. Each bar represents an individual CpG, and represents the ratio of methylated to unmethylated reads, similar to (E). (H) Quantification of gMFI of E2F2 (p-value= 0.0111) protein expression on CD8+ T cells in the blood from Aza or PBS-treated mice. Data is pooled from 2-4 independent experiments with n=8-14 mice per group, as shown. Statistical analyses were performed using Mann-Whitney U test, with p-values as shown.

We identified thousands of differentially methylated regions (DMRs) in both T cell subsets with Aza treatment **(Supplemental figure 4A)**. The total number of DMRs as well as the relative distribution of hypermethylated vs hypomethylated regions, was comparable across both T cell subsets **(Supplemental figure 4B)**. After annotating the genes nearby or overlapping with DMRs, we manually curated a set of differentially methylated immune genes that have an established role in T cell biology **(Figure 4 B-C)**. This selection was done to prioritize biologically relevant epigenetic changes, and the complete list of DMRs is available in supplemental data. Since exhaustion is well-characterized in CD8+ T cells, we then decided to focus on that population for further analysis.

To get a deeper understanding of the functional relevance of these changes, we performed pathway enrichment analysis on all the DMRs using GREAT^41,42^. Among the top enriched pathways was the β-Catenin/TCF signaling axis **(Figure 4D)**. This pathway is notable given its established role in maintaining stem-like and memory T cell states^45,46^. In particular, the transcription factor TCF1, encoded by *TCF7*, is a critical mediator of stem-like/progenitor exhausted and memory biased T cell programs, and is and is known to be epigenetically regulated in exhaustion settings^15–17,47^.

Consistent with these findings, we observed hypomethylation at promoter regions of *TCF7* in CD8+ T cells upon Aza treatment **(Figure 4B & E)**. Flow cytometry analysis further revealed increased expression of TCF1 protein in circulating CD8+ T cells from Aza-treated mice **(Figure 4F)** but not in spleens **(Supplemental figure 4C)**. This data suggests that Aza promotes hypomethylation of key loci associated with stem-like T cell programs, which can potentially restore T cell function.

### Aza increases expression of E2F2 in CD8+ T cells

In addition to TCF1, the E2F family of transcription factors are also part of the β-Catenin/TCF axis. Although these transcription factors are classically characterized for their role in cell proliferation and cell cycle progression, emerging evidence suggests potential crosstalk between E2F signaling and T cell differentiation programs^48–50^. Given recent reports suggesting that E2F family member, E2F1 can directly regulate *TCF7* expression^51^, we focused on E2F2 from our screen, which has known functional redundancy with E2F1. RRBS analysis revealed hypomethylation of E2F2 in CD8+ T cells upon Aza treatment **(Figure 4B & G)**. We also observed a significant increase in E2F2 protein expression in CD8+ T cells in the blood of Aza-treated mice compared to vehicle **(Figure 4H),** but not in spleens (**Supplemental figure 4D)**. While the exact mechanism by which E2F2 and TCF1 interact in our model remains to be defined, these findings highlight that Aza-induced epigenetic remodeling may involve a broader network of transcription factors which regulates differentiation states and proliferation.

### Aza-induced phenotypic changes are preserved in murine *ex vivo* cultures

To determine whether the immune phenotypes induced by Aza *in vivo* could be recapitulated in a controlled setting, we next evaluated the effects of Aza on murine T cells *ex vivo*. Splenocytes were isolated from leukemic mice and were cultured for five days in the presence of PBS or two doses of Aza (500nM or 1μM) **(Figure 5A)** and subsequently analyzed for relevant T cell markers.

**Figure 5:**
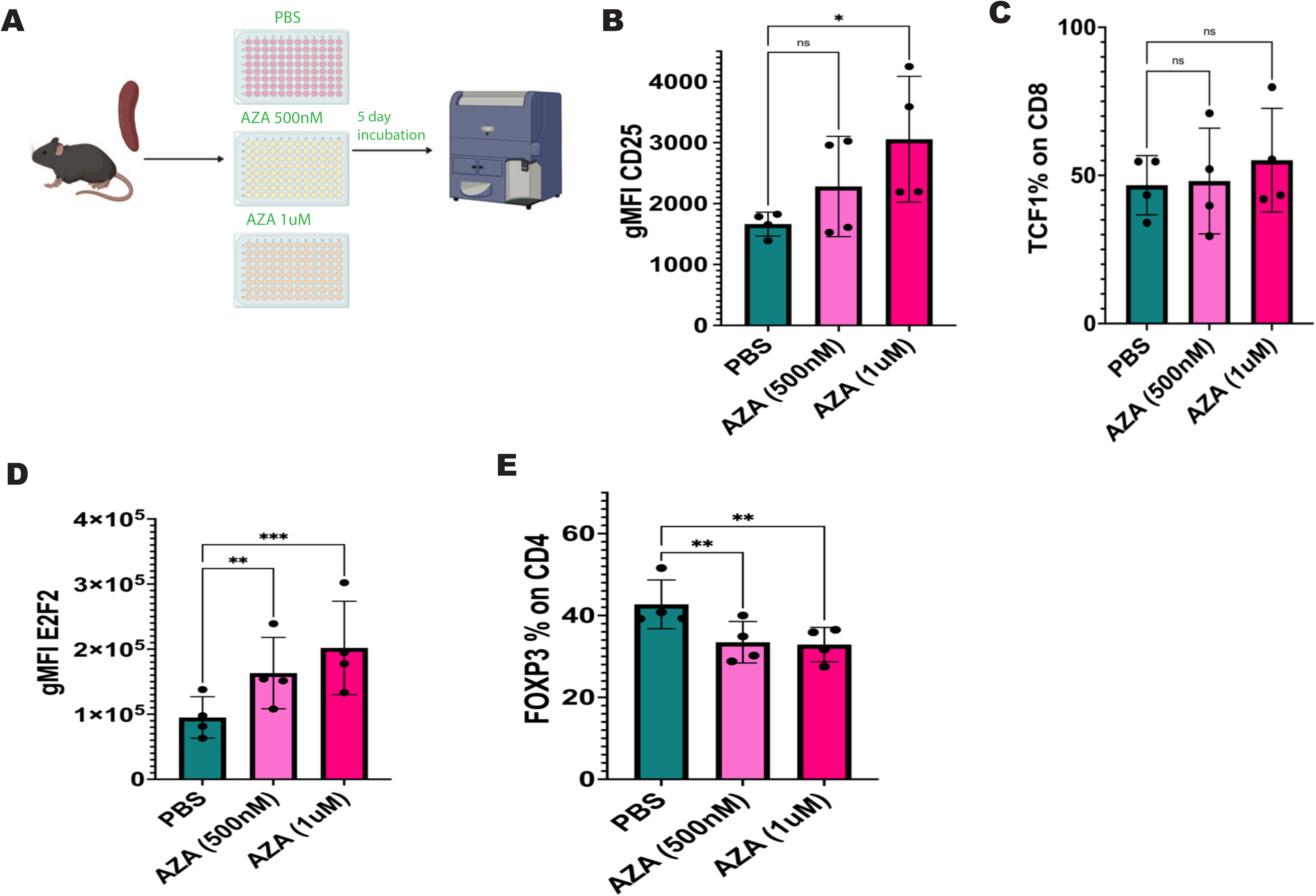
Aza-induced changes in murine *ex vivo* cultures. (A) Schematic of the *ex vivo* experimental design. Splenocytes were isolated from leukemic mice and cultured for 5 days. Cultures were treated daily with PBS or Aza at 500nM or 1μM. At the end of the culture period, cells were harvested for flow cytometric analysis for T cell phenotyping. (B) Quantification of gMFI of CD25 expression on CD8+ T cells comparing PBS vs 500nM Aza, (p-value= 0.1869) and 1μM Aza (p-value= 0.0101). (C) Frequency of TCF1+ cells among CD8+ T cells comparing PBS vs 500nM Aza (p-value= 0.869), and 1µM Aza (p-value= 0.562). (D) Quantification of gMFI of E2F2 expression in CD8+ T cells comparing PBS vs 500nM Aza, (p-value= 0.0035) and 1μM Aza (p-value= 0.0007). (E) Frequency of FOXP3+ regulatory T cells among CD4+ T cells comparing PBS vs 500nM Aza (p-value= 0.0051) and 1μM Aza (p-value= 0.0051). Data is pooled from two independent experiments with n=2 mice per experiment. Statistical analysis was performed using repeated measures one-way ANOVA.

Consistent with our *in vivo* observations, Aza treatment led to a dose-dependent immune modulation, with the 1μM dose eliciting the most pronounced effects. Specifically, Aza increased CD25 expression and E2F2 levels in CD8+ T cells **(Figure 5B & D)**, while TCF1 was not significantly altered **(Figure 5C)**. We also saw a significant decrease in the frequency of FOXP3+ Tregs **(Figure 5E)**. Together, these data indicate that key Aza-induced immune features observed *in vivo*, particularly enhanced T cell activation and transcriptional remodeling are preserved *ex vivo*. This data recapitulates key *in vivo* features of Aza-mediated immune reprogramming, supporting a T cell–intrinsic component within a controlled microenvironment.

### Aza modulates T cell activation and phenotype in human AML samples

We evaluated the impact of Aza treatment on T cells in samples from patients with AML. Frozen PBMCs with diverse mutation profiles **(Table 3)** were cultured *ex vivo* on low dose IgG or anti-CD3 coated plates and treated with vehicle or Aza (500nM) for 5 days **(Figure 6A)**. Cultures were supplemented with cytokines to support both leukemic blasts (IL-3, IL-6 and SCF) and T cell survival (IL-2, IL-7 and IL-15).

**Figure 6:**
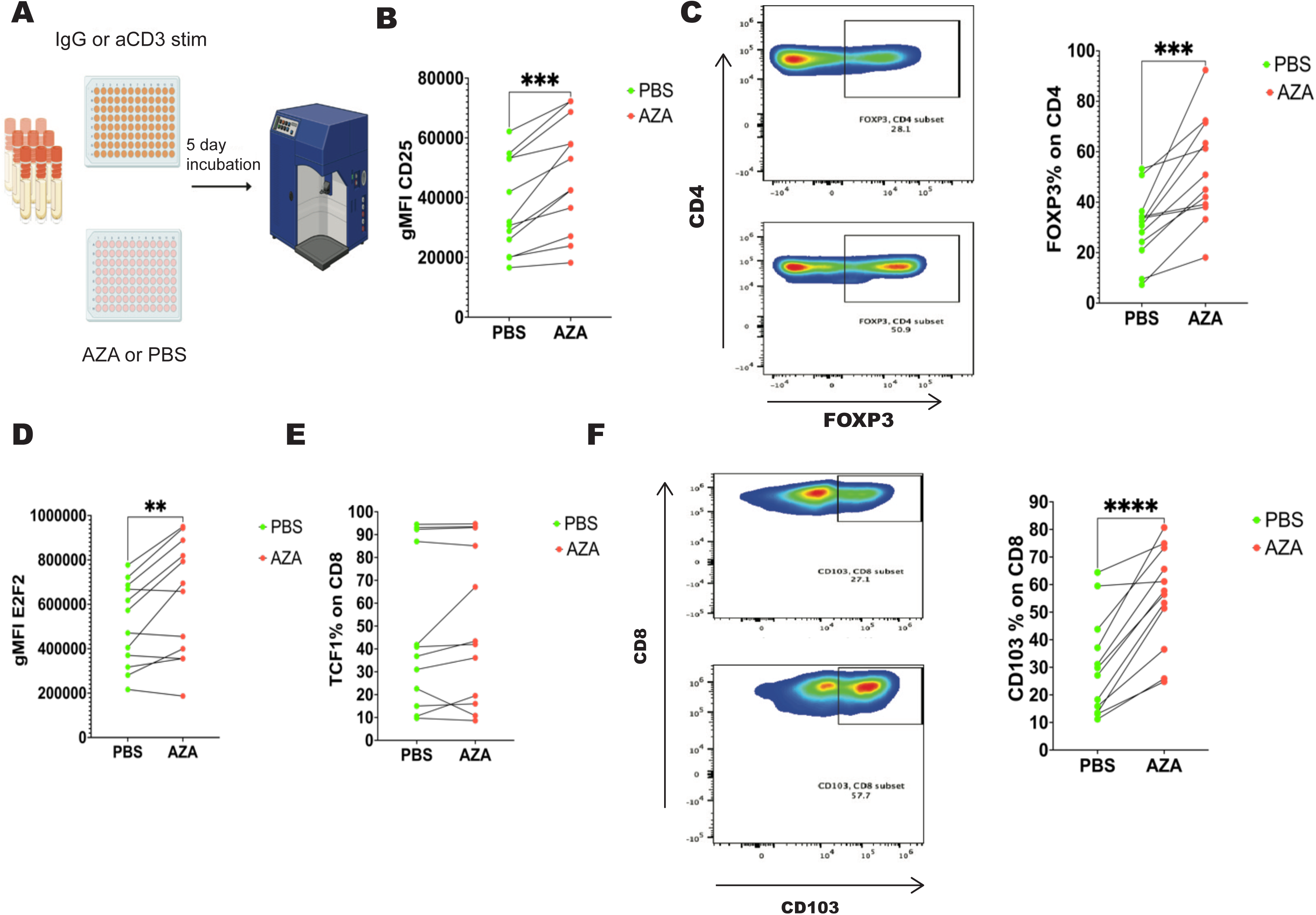
Aza-induced changes in *ex vivo* cultures of AML patient samples. (A) Experimental schematic of the *ex vivo* culture assay using primary human samples with AML. Cryopreserved peripheral blood mononuclear cells (PBMCs) from patients with AML having diverse mutation profiles (Table 3) were plated on IgG or anti-CD3-coated plates. Cells were cultured in cytokine supplemented media and treated daily with Aza or PBS for five days prior to flow cytometric analysis. (B) Quantification of gMFI of CD25 expression on CD8+ T cells following anti-CD3 stimulation in PBS vs Aza-treated samples. Each data point represents an individual patient sample and each line connects paired samples from the same patient. (p-value= 0.0002). (C) Representative dot plots depicting FOXP3 expression on CD4+ T cells (left), and corresponding quantification (right) of the frequency of FOXP3+ cells among CD4+ T cells (p-value= 0.0004). (D) Quantification of gMFI of E2F2 expression measured among CD8+ T cells with PBS or Aza treatment. p-value= 0.0052. (E) Frequency of TCF1+ cells among CD8+ T cells (p-value= 0.276). (F) Representative dot plots depicting CD103 expression on CD8+ T cells (left) and corresponding quantification (right) of the frequency of CD103+ among CD8+ T cells (p-value= <0.0001). Statistical analyses were performed using a paired t-test, and a p-value of <0.05 was used to determine significance. n=12 patient samples.

**Table 3:** Overview of patient samples characteristics-.

| Patient ID | Gender | Mutations | Diagnosis |
| --- | --- | --- | --- |
| Patient 1 | Female | NPM1, RAD21, DNAJC21, **DNMT3A**, PTPN11 | Initial diagnosis |
| Patient 2 | Male | NPM1, ZBTB7A, RTEL1, KMT2D, **DNMT3A** | Initial diagnosis |
| Patient 3 | Male | ZRSR2, **TET2**, EZH2, CBL, ASXL1, CUX1, DOCK8, | Post chemotherapy |
| Patient 4 | Male | IGLL5, NRAS, USH2A, ASXL2, CASP10, BCORL1, U2AF1 | Initial diagnosis |
| Patient 5 | Female | FLT3-ITD, ATM, TET2, SETD2, NPM1, RYR1, KDM6A | Relapse |
| Patient 6 | Male | NRAS, NF1, ASXL1, WAS, STAG2, EED, **TET2**, CD27, RUNX1 | Post chemotherapy |
| Patient 7 | Female | NPM1, PHF6, PRDM1 | Initial diagnosis |
| Patient 8 | Male | ASXL1, RUNX1, CSF3R, NRAS, BRAF, FLT3, EZH2, CREBBP, CBL | Post chemotherapy |
| Patient 9 | Male | FLT3-ITD, NPM1, CHEK2, **TET2**, DNMT3A, ARID1B, FAT1, GNA13, SPEN | Initial diagnosis |
| Patient 10 | Female | BCOR, FAT4, NRAS, SPEN | Initial diagnosis |
| Patient 11 | Male | KDM6A, CBL | Initial diagnosis |
| Patient 12 | Female | FLT3-ITD, **DNMT3A**, ATG2B, RUNX1 | Initial diagnosis |

We observed that IgG stimulation did not induce appreciable T cell activation or phenotypic changes under either treatment condition. Samples that were incubated with anti-CD3 had a higher T cell viability allowing us to assess several T cell markers. Under these conditions, Aza treatment led to a significant increase in CD25 expression on CD8+ T cells compared to vehicle-treated samples **(Figure 6B)**, indicating enhanced T cell activation. Interestingly, Aza treatment led to an increase in FOXP3+ Tregs in our patient samples, which is consistent with prior literature^52,53^ **(Figure 6C)**. We also examined the expression of transcription factors identified in our murine epigenetic analysis. Aza treatment led to a significant increase in E2F2 expression in CD8+ T cells across patient samples **(Figure 6D)**. In contrast to the *in vivo* treatment in the mouse model, we did not observe a significant increase in TCF1 following Aza treatment of human samples **(Figure 6E)**. This difference may reflect the shorter duration of Aza treatment or microenvironment differences observed *ex vivo* vs *in vivo*. Additionally, Aza caused an increased frequency of CD103+ CD8+ T cells **(Figure 6F)**, indicating a shift towards a resident memory population.

Overall, this data demonstrates that Aza can modulate activation and transcriptional machinery in human CD8+ T cells in the context of AML.

## Discussion

Immunotherapy has transformed treatment across several hematologic malignancies, especially B cell malignancies. However, therapies that have worked well in lymphoid malignancies like immune checkpoint blockade, T cell-redirecting antibodies and CAR T cells have shown low efficacy in AML^54–58^. We believe that immune suppression including T cell exhaustion is a major barrier to effective immunotherapy in AML. In support of that, patients who relapse with AML exhibit more severe exhaustion phenotypes of their T cells compared to newly diagnosed patients^9,59,60^. Epigenetic remodeling is a central driver of T cell exhaustion, and the Youngblood group showed that DNMT3a deletion in T cells restores effector function in a chronic LCMV infection model, highlighting DNA methylation as a tractable mechanism for reversing exhaustion^21^.

Aza was originally developed as a cytotoxic agent. It was later realized that it inhibits DNA methyltransferases, leading to global hypomethylation^61–64^. This property has been linked to effects in tumor-intrinsic properties, including reactivation of tumor suppressor genes and inhibition of leukemia proliferation. More recently, however, accumulating evidence suggests that Aza also exerts tumor-extrinsic immune modulatory effects. In both solid tumors and AML, Aza has been shown to increase expression of cancer testes antigen, enhance antigen presentation by upregulating MHC expression and induce type I interferon signaling via expression of endogenous retroviruses in cancers^29–31,65–68^.

Despite these advances, relatively little is known on how Aza directly impacts T cell exhaustion and its epigenetic reprogramming in AML. In this study, we demonstrate that Aza treatment reduces leukemic burden in a spontaneous genetically-engineered mouse model of AML. Importantly, these changes are not limited to tumor reduction but are accompanied by functional and phenotypic restoration of exhausted T cells. Treatment restores T cell proliferative capacity, reshaping memory differentiation and induces epigenetic remodeling in T cells.

A recent phase III clinical trial using oral Aza as maintenance therapy for AML following intensive chemotherapy reported immune modulation in patients post remission^69^. In fact, Aza treatment led to a decrease in expression of exhaustion markers PD-1 and TIM-3 in T cells. In contrast, upregulation of immune inhibitory checkpoint markers including PD-L1 and PD-L2 was observed in leukemia cells.

Clinically, although Aza therapy results in meaningful responses in a subset of patients with AML, less than half of the patients respond, and more than 50% of responders relapse within two years^28^. Combination strategies targeting immune checkpoints, particularly PD1/PDL1, have been proposed as a means to overcome resistance^70,71^. However, recent clinical trials combining Aza with a PD-1 blockade demonstrated non-durable responses^72^. Our findings suggest that this lack of efficacy can be due to the presence of additional, PD-1-independent, mechanisms of immune resistance in AML. While Aza restored T cell proliferation in most of the mice with AML, some mice failed to restore proliferation, suggesting that these changes alone may be insufficient to overcome the complex immune suppressive landscape in AML.

Prior work from our group demonstrated that epigenetic reader inhibition via BET inhibitors (BETi) can restore T cell function in our AML mouse model^32^. BETi treatment increased expression of precursor exhausted T cell genes including TCF1, SLAMF-6 and CXCR5, enhanced chromatin accessibility at progenitor program loci, and restored T cell proliferation in combination with checkpoint blockade. While BETi primarily work by altering transcription through chromatin acetylation, Aza targets DNA methylation, which is more stable. Despite these mechanistic differences, both approaches restore the stem-like progenitor states in T cells, suggesting that multiple epigenetic mechanisms may work together to enforce exhaustion in AML. Targeting complementary epigenetic mechanisms may therefore provide synergistic benefit in restoring anti-tumor T cell effects in AML.

While patients with AML have been reported to exhibit increased Treg frequencies^73–75^, and Aza treatment further increased FOXP3+ Tregs in human samples^76,77^, we consistently observed a reduction in Tregs in Aza-treated mice with AML. This decrease was also observed when murine samples were treated *ex vivo*. This is likely because murine T cells in leukemic hosts are chronically activated in an antigen-experienced environment, which may make Tregs more susceptible to hypomethylation-induced effects. In contrast, human PBMC-based cultures may preferentially reveal direct epigenetic impact of Aza on FOXP3+ Tregs in the absence of microenvironmental constraints^52,53^, highlighting that the tumor microenvironment plays a critical role in determining the effects of hypomethylating agents.

Overall, our work provides the mechanistic framework for rational combination studies and highlight the promise of targeting both tumor intrinsic and immune pathways to achieve durable responses in AML.

## Supporting information

Sig DMR table

## Acknowledgements

This work and EFL was supported by NCI 5R01CA262145 (Lind PI), ARTNet 5U54CA224019 (Tyner PI). RP was supported in part by Integrated Cancer Biology Stipend Support Award provided by OHSU. We would like to thank the Flow Cytometry Core (RRID: SCR_009974), Advanced Light Microscopy Core (RRID: SCR_009961), the KCVI Epigenetics consortium, the Advanced Computing Center, and the Massively Parallel Sequencing Shared Resource (MPSSR) Core (RRID: SCR_009984). We would also like to thank Dr. Stefanie Kaech-Petrie for image analysis, and Kandace Wheeler for preparing RRBS libraries.

## Authorship Contributions

RP and EFL contributed to conceptualization, study design, investigation, and writing. RP, YK, and JM performed the experiments. RP conducted data collection and analysis. CEL performed quality controls and cleaned and concentrated DNA samples for RRBS. BAD and LC performed RRBS data analysis. All authors edited the manuscript.

## Supplemental figure legends

**Supplemental figure 1:**
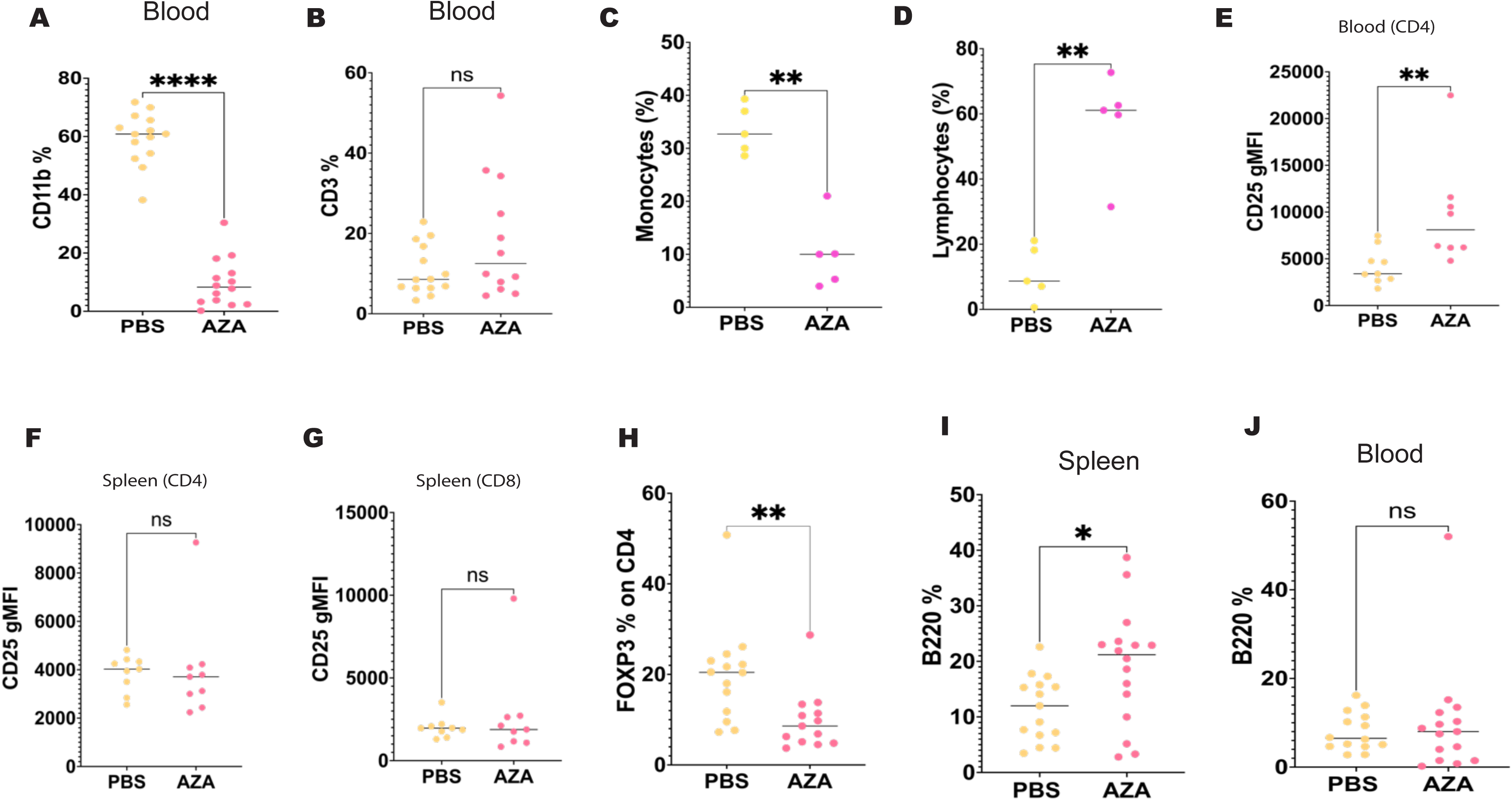
Aza’s impact on tumor burden in AML mouse model. (A-B) Quantification of the frequency of CD11b+ myeloid cells (A) and CD3+ T cells (B) among live cells in peripheral blood of AML mice treated for three weeks with Aza or PBS, as assessed by flow cytometry. p-value= <0.0001 (A), 0.252 (B). (C) Frequency of monocytes quantified using automated hematology analyzer at the study end point, comparing Aza and PBS treated mice. p-value= 0.0079. (D) Frequency of lymphocytes quantified using automated hematology analyzer at the study endpoint, comparing Aza- and PBS-treated mice. p-value= 0.0079. (E) Geometric mean fluorescence intensity (gMFI) of CD25 expression on CD4+ T cells, measured in the blood after three weeks of PBS or Aza treatment. p-value= 0.0055. (F) (G) gMFI of CD25 expression on CD4+ T cells (F) (p-value= 0.341) and CD8+ T cells (G) (p-value= 0.863) isolated from spleens of PBS and Aza treated mice after three weeks of PBS or Aza treatment. (H) Frequency of FOXP3+ cells in CD4+ cells, in blood of Aza- and PBS-treated AML mice. p-value= 0.0033. (I-J) Frequency of B cells, identified by B220 expression among live cells in the spleen (I) (p-value= 0.0220) and blood (J) (p-value= 0.747), respectively, of AML mice treated for three weeks with Aza or PBS. Data was analyzed using GraphPad Prism. Statistical significance determined using Mann-Whitney U test. Each data point represents an individual mouse.

**Supplemental figure 2:**
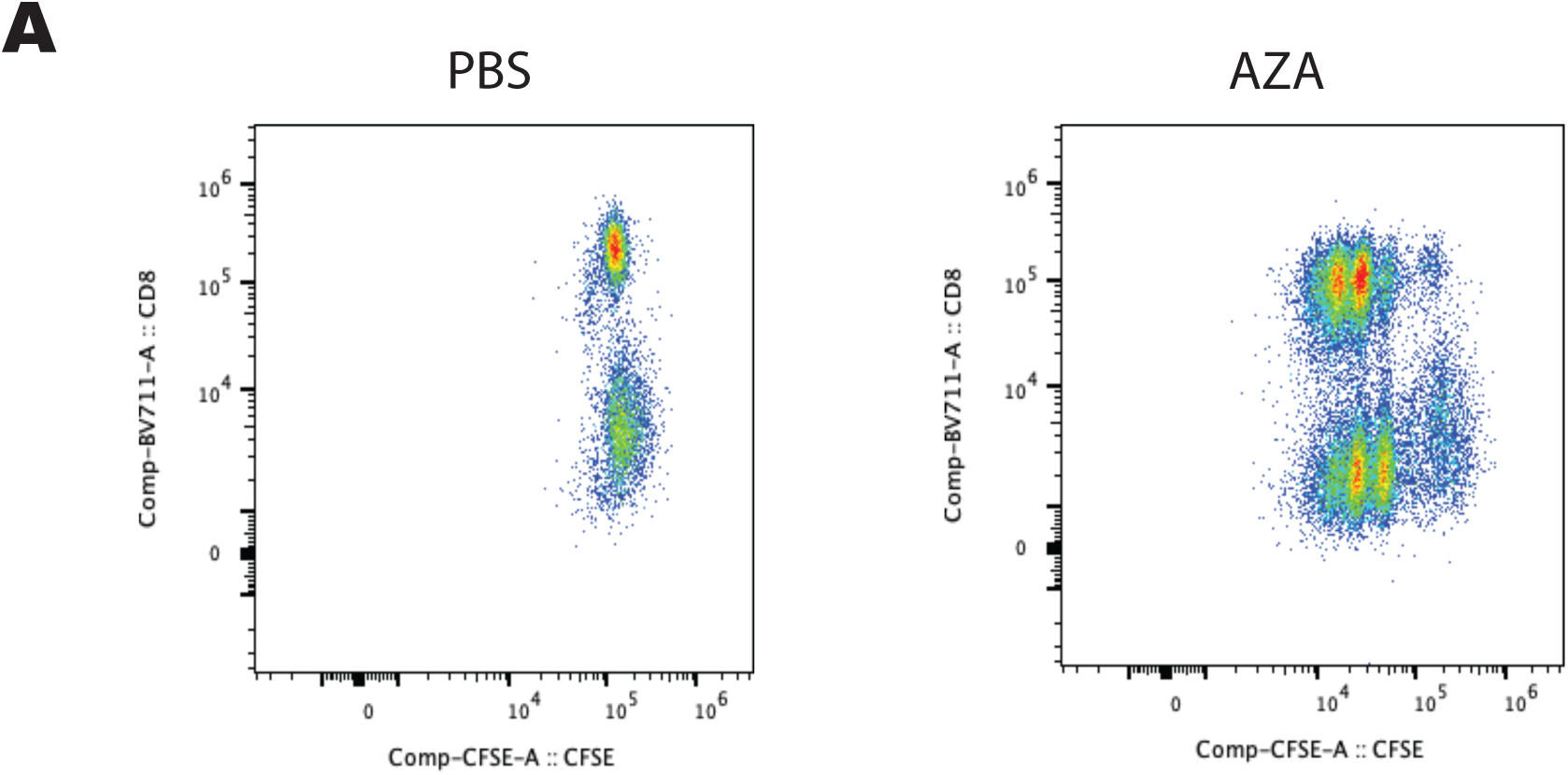
Proliferation post Aza treatment. (A) Representative flow cytometry dot plots comparing PBS and Aza treated mice following anti-CD3 stimulation, showing CFSE (x-axis) vs CD8 (y-axis). Plots are gated on live CD3+ T cells.

**Supplemental figure 3:**
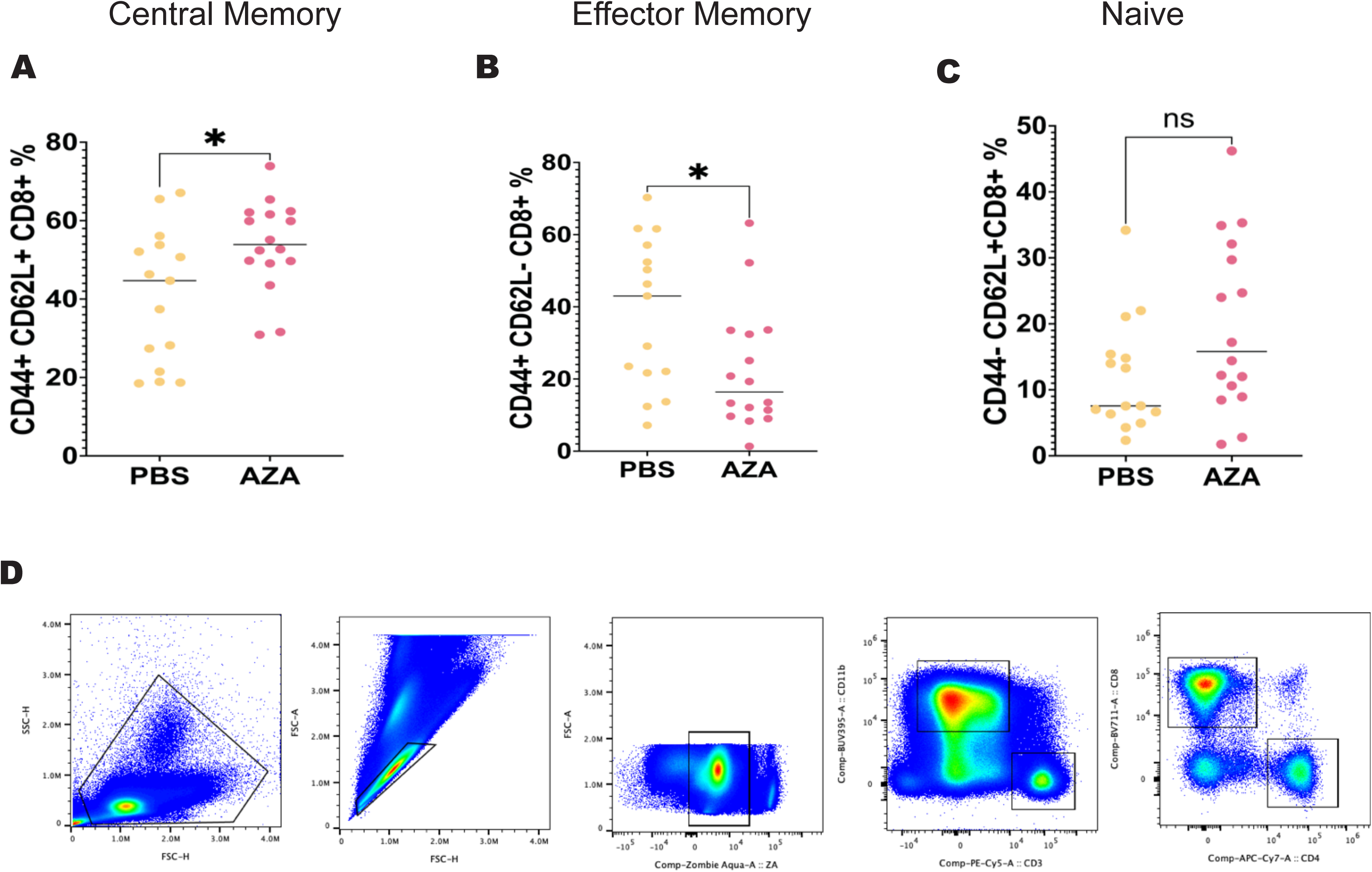
Memory subset changes after Aza treatment in spleens of AML mice. (A-C) Quantification of central memory (CD44+ CD62L+; A), effector memory (CD44+ CD62L-; B) and naïve (CD44-CD62L+; C) CD8+ T cell subsets in the spleens of AML mice treated with PBS or Aza for three weeks. p-value for central memory is 0.0357, effector memory is 0.033, naïve is 0.0856. (D) Representative gating strategy used for analyzing flow cytometry data using FlowJo. Data is pooled from 4 independent experiments. Statistical significance was determined using Mann-Whitney U test. Each data point represents an individual mouse.

**Supplemental figure 4:**
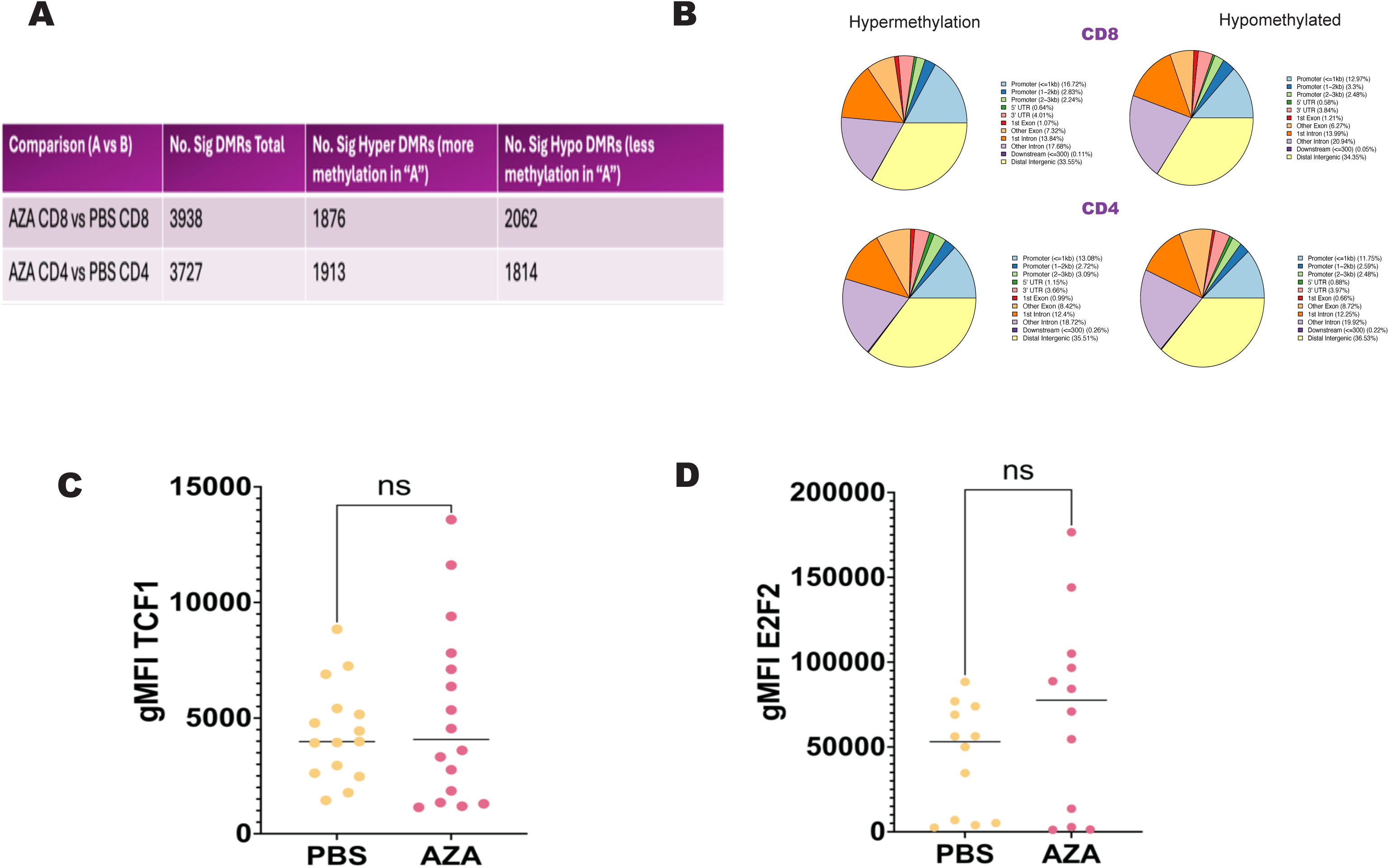
RRBS revealing hypomethylation changes as well as protein expression changes in spleens post treatment. (A) Table summarizing the total number of differentially methylated regions (DMRs) identified in CD4+ and CD8+ T cells of Aza or PBS-treated mice. (B) Pie charts showing the genomic annotation of differentially methylated regions (DMRs) identified in CD8+ (top) and CD4+ (bottom) T cells, categorized by hypermethylated (left) and hypomethylated (right) regions across treatment groups. The DMRs were distributed across multiple genomic regions, including promoters, gene bodies and intergenic regions, consistent with widespread epigenetic remodeling. (C-D) Geometric mean fluorescence intensity (gMFI) of TCF1 expression (C) (p-value= >0.999) and E2F2 (D) (p-value= 0.2913) measured by flow cytometry in splenic CD8+ T cells from leukemic mice following three weeks of Aza or PBS treatment *in vivo*. Cells were gated on live CD3+ CD8+ T cells prior to analysis. Data is pooled from 3-4 independent experiments. Statistical significance was determined using Mann-Whitney U test. Each data point represents an individual mouse.

